# Disambiguating brain functional connectivity

**DOI:** 10.1101/103002

**Authors:** Eugene P. Duff, Tamar Makin, Stephen M. Smith, Mark W. Woolrich

**Affiliations:** FMRIB Centre, University of Oxford, Oxford, United Kingdom, OX3 9DU; Oxford Centre of Human Brain Activity, University of Oxford, Oxford, United Kingdom, OX3 7JX

**Keywords:** Correlation, SNR, functional connectivity, effective connectivity, FMRI

## Abstract

Functional connectivity (FC) analyses of correlations of neural activity are used extensively in neuroimaging and electrophysiology to gain insights into neural interactions. However, correlation fails to distinguish sources as different as changes in neural signal amplitudes or noise levels. This ambiguity substantially diminishes the value of FC for inferring system properties and clinical states. Network modelling approaches may avoid ambiguities, but require specific assumptions. We present an enhancement to FC analysis with improved specificity of inferences, minimal assumptions and no reduction in flexibility. The Additive Signal Change (ASC) approach characterises FC changes into certain prevalent classes involving additions of signal. With FMRI data, the approach reveals a rich diversity of signal changes underlying measured changes in FC, bringing into question standard interpretations. The ASC method can also be used to disambiguate other measures of dependency, such as regression and coherence, providing a flexible tool for the analysis of neural data.

---

Correlation and regression are widely used to characterise the extent to which sets of signals are related, and how these relations might change over time or across experimental conditions. For example, in neuroscience, functional connectivity (FC) analyses use correlation or related measures to identify networks showing shared activity and to characterise changes in these relationships over time (1–5). While these analyses lend themselves to interpretation in terms of intrinsic connectivity of brain regions, correlation will be sensitive to a wide variety of changes in signal dynamics (1–3, 6, 7). For example, differences in noise levels will alter correlation, as will changes in the amplitude of shared neural activity, or changes in higher order properties of constituent signals (1–3). Similar challenges occur in the frequency domain, where analogues of correlation such as coherence can be influenced by a variety of very different changes in signal and noise properties (3). This diversity of potential phenomena producing changes in correlation makes the interpretation of reported changes in functional connectivity studies highly uncertain. These ambiguities reduce the usefulness of correlation-based approaches for characterising network structure and for guiding the specification and interpretation of more complex, multivariate modelling approaches that aim to distinguish directed and mediated relationships between nodes (1, 4, 8, 9).

Various approaches posit specific models of signal to avoid the ambiguities of simple correlational analyses. Approaches include partial correlation (4, 9), structural equation modelling (SEM) (8), dynamic causal modelling (DCM) (10, 11), Granger causality(12) and the distribution-based causal inference approach, LiNGAM (13). A variety of information may inform the estimation of these models, including the stationary or dynamic covariance structure, higher order moments, and timing discrepancies between regions. If appropriately defined, these approaches may avoid sensitivity to signal-to-noise changes through their ability to model multiple sources of variance in the network. However, care is required to ensure that model assumptions are realistic, and the various signal and noise sources are appropriately modelled. In neuroscience, there will be various neural and physiological processes affecting signals, across a range of spatial and temporal scales. Signal sources (e.g. measurements from distinct spatial regions) must be appropriately selected - mis-specified, missing, or redundant nodes can have a significant effect on network identification and estimation (14). Network modelling approaches such as Granger, SEM and DCM are designed to reveal causal structure, which may not provide optimal characterisation if causality cannot be reliably estimated, for example, when redundant nodes providing common signal are present. If complex network models do not appropriately account for major aspects of dynamics, missed effects can influence available network parameters in complex ways. Simple, flexible measures of similarity that make minimal assumptions are important complementary approaches to provide insight into those aspects of dynamics that need to be modelled. Here we present a complementary approach which directly enhances correlation analysis to provide more specific insight into underlying dynamics. With minimal additional assumptions, the approach identifies additive signal changes (ASC), a class of changes that is intended to encompass a variety of common changes in many systems. This class, for example, covers changes in noise levels, or the amplitude of a signal driving correlation between two nodes. A feature of this class of changes is that correlation changes are always accompanied by changes in variance. Variance changes have been often identified in parallel to correlation, but not regularly characterised alongside them (2, 6, 15, 16). By tracking variance, inferences can be made regarding whether observed changes in correlations might be explained by specific types of additions of signal. The approach is related to the network-inference approach of structural equation modelling and related approaches such as dynamic causal modelling, but here the focus is on deriving a basic bivariate analysis that is computationally simple, and permits the mapping of connectivity mapping and rapid exploratory analyses. Cole et al focus on similar phenomena, outlining a conjunction-based approach that combines correlation and covariance to provide insight into possible shared-signal based dynamics underlying FC changes (3). This approach can be valuable for exploring the sources of FC. However, it does not provide inferences directly relating to putative underlying signal changes.

Our enhanced ASC correlation analysis strategy can be used to: 1. Produce more interpretable seed based, independent component analysis (ICA), network matrix and psycho-physiological interaction (PPI) FC analyses. 2. Make inferences regarding the networks in which signals are embedded. 3. Appraise network modelling by identifying the distinct types of changes in data that underlie network model fits. In the following, we describe the approach and inference procedures, including extensions to partial correlation and coherence analysis. We then test the approach on simulated and empirical datasets. The results provide insight into the extent to which functional connectivity and related neuroscientific analyses can be driven by simple additive signals. We also describe in detail the relationship of this approach to network modelling.

## Results

### Model

We are interested in making inferences regarding the source of observed changes in correlation of pairs of signal sources across different (e.g. cognitive) conditions. Consider a pair of nodes (e.g. brain regions) ***X*** and ***Y*** which can be measured in conditions ***A*** or ***B***. We consider the node signals during a specific condition (e.g. ***X***_***A***_) as being produced by an ergodic stochastic process providing a covariance matrix ∑_*X_A_*, *Y_A_*_:

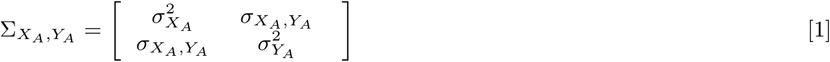

Where 
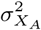
 is the variance of node ***X*** in condition ***A***, and σ_*X_A_*, *Y_A_*_ is the covariance between nodes. Estimates from data of how these riances change across conditions (**Q_A_**, **Q_B_**) will be used for inferences about differences between underlying processes across the conditions.

***Defining additive changes in signal*.** Our analysis approach focuses on determining whether differences in the correlation between nodes ***X*** and ***Y*** across conditions ***A*** and ***B*** can be explained by certain additions of new signal into one or both of the nodes. To define signal additions, we first posit a baseline stochastic process present in an individual node in both conditions, e.g. for node **X**, **S_X_**. The additive class corresponds to differences in correlation that are produced by an addition of a further stochastic process, **N_X_** (or similarly **N_Y_**) in one of the two states e.g.:

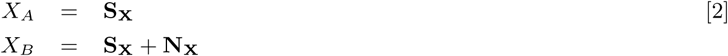

Distinct additions may occur in both nodes, and may appear in condition *A* for one node, and *B* in the second, thus modelling both increases and decreases in signal relative to condition **A**. We place one constraint on the new signal: **N_X_** cannot be negatively correlated with **S_X_** (*ρs_X_*, *N_X_* ≥ 0). This ensures that the new signal is additive, and not negating the variance associated with 
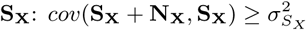
.

The additive class of signal change covers a variety of basic changes in signal properties that can be expected to occur in many contexts (Fig. 1.1-1.3). The new signal **N_X_** may correspond to a process that is already a component of the baseline signal (therefore reflecting an increase in the amplitude of this component in one condition), or it could correspond to the introduction of novel signal. The addition of a process that is uncorrelated to activity in the second node will reduce the correlation between nodes. Such a change could reflect an increase in uncorrelated noise levels. Conversely, the addition of some common process into both nodes will increase correlation. The additive class does not restrict the correlation of **N_X_** and **N_Y_**, and so includes scenarios where these additional signals overlap. As the additive signal can appear in either condition, the class encompasses both increases and decreases in signal, and changes where signal additions occur in different conditions for each node.

**Fig. 1.**
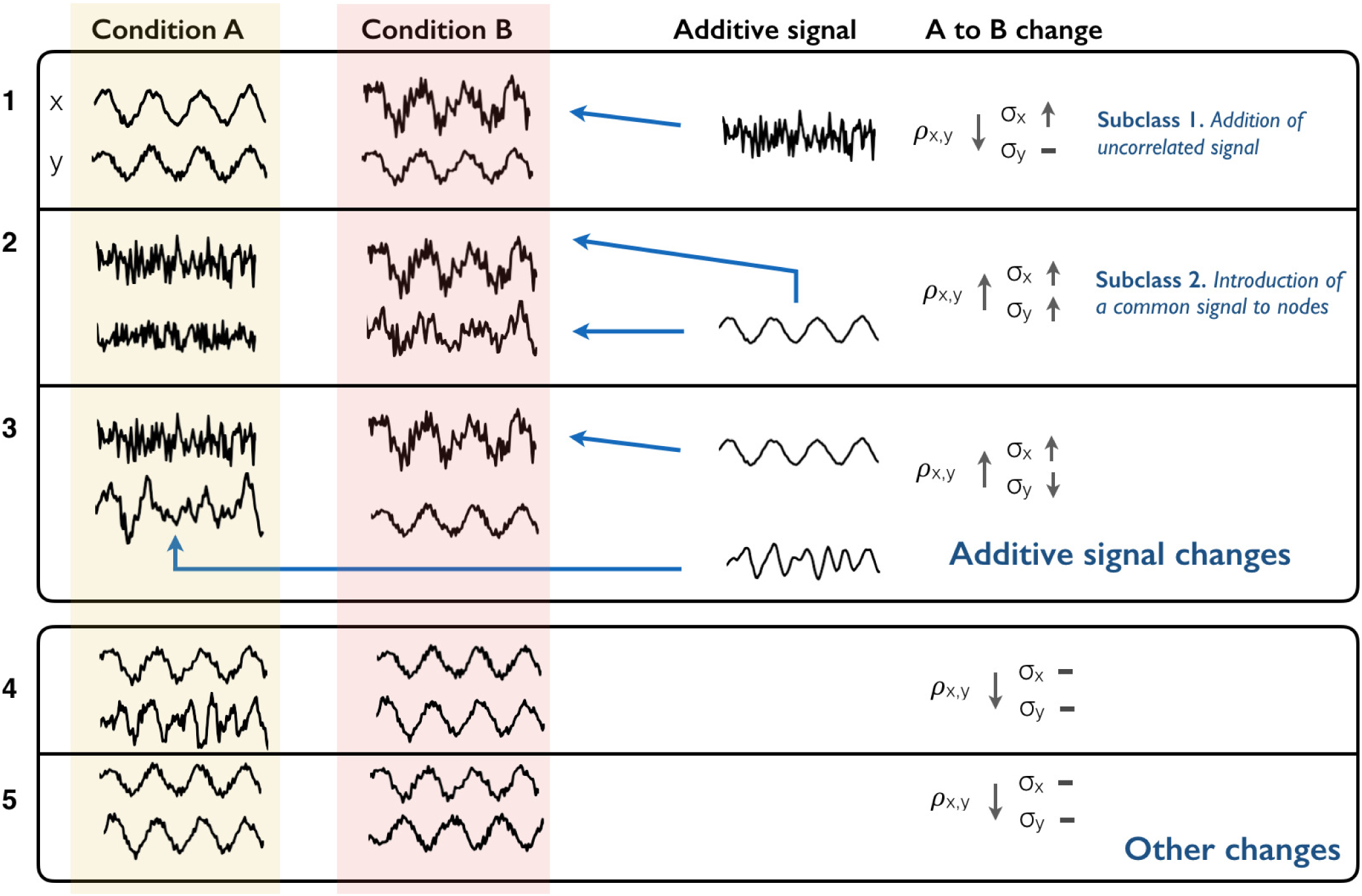
Representation of effects of additive signal changes (ASC) on correlation. The upper box demonstrates three examples of additive signal changes to correlation. The blue arrows represent the addition of signal into a node in a certain condition. The first example corresponds to the subclass of additions in uncorrelated signal. Here, signal uncorrelated with region **Y** is added to **X** in condition *B*, reducing correlation and increasing variance. In the second, a common signal is added to both regions in condition *B*, increasing correlation and variance. In the third example, region **X** receives an addition of signal already present in region **Y**. At the same time, some signal not shared by region **X** (e.g. some input from a third region) is removed from region Y in condition *B*. The overall effect is an increase in correlation. The second box shows some scenarios that do not fall within the class of additive changes. The first example shows a synchronization of signals whose temporal properties, including variance, otherwise do not substantially change. The final example shows two signals where their correlation flips from positive to negative. This could be explained by the addition of a great deal of negatively correlated signal, but falls outside our definition of additive signal.

A variety of scenarios do not fall into the class of additive changes. In terms of the model used here, such changes correspond to situations where the baseline signal is replaced, modified, or changes sign. A second condition may involve a mixture of increases and decreases in the amplitude of existing signals. For example two signals may purely synchronize, with no change in variance.

***Variance changes with ASC*.** A characteristic feature of additive signal changes is that the variance of at least one node will change whenever there is a change in correlation, with nodes having a higher variance in the condition where they receive the additional signal. Larger changes in correlation will typically produce larger changes in variance, with the specific change being dependent on the initial correlation and the nature of the new signal. Given observed variance changes, it is possible to determine whether particular changes in FC can be explained by ASC by calculating bounds on *ρ_X_B__*,*_Y_B__* across the range of possible new signals, *N_Y_* and *N_X_*:

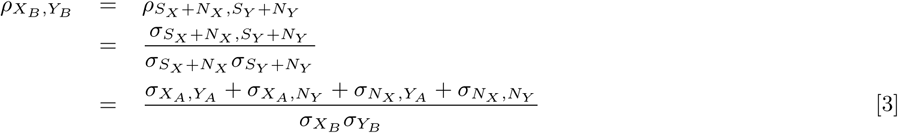

Correlation can be expressed in terms of observables and *ρX_A_*,*N_X_* and *ρY_A_*,*N_X_*, by obtaining expressions the remaining parameters using relationships derived from expressions for observed variance (see Materials and Methods). This permits the identification of the range of possible changes in correlation that could be explained by ASC. An approach to accommodate observation error is described in Statistical Inference, below.

***Subtypes of Additive changes in signal*.** A variety of distinct additions of signal are covered by the ASC class. It is possible to define subtypes of ASCs and to test for evidence that they specifically can explain observed changes:

*1. Addition of uncorrelated signal*. This subclass covers scenarios such as changes in levels of uncorrelated measurement noise, and changes in the amplitude of signal components that are independent of activity in the second node. The additional signals are uncorrelated with signal in the other node. That is, **N_X_** is uncorrelated with both **S_Y_** and **N_Y_**, and **N_Y_** is uncorrelated with **S_X_** and **N_X_** (e.g. Fig. 1.1). If the uncorrelated signal is added in condition *B*, then Eqn. 4 becomes:

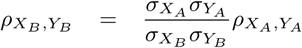

For this scenario, a given change in variance will predict a specific change in correlation (excluding observational uncertainties), with increases in correlations predominately accompanied by decreases in variance.

*2. Introduction of a common signal to both nodes.* This class corresponds to the scenario where there is an addition of the same latent signal component to both nodes, i.e., *ρ_N_X__*,*_N_Y__* = 1 (Fig. 1.2). The amount of this addition is be determined by the observed changes in variance. Here, in Eq. 2, we set **N_X_** = *s_N_X__***N_c_**, **N_y_** = *s_N_Y__***N_c_**, where **N_c_** is a latent stochastic process set to have variance of 1, and *s_N_X__* and *s_N_Y__* are scaling factors, such that *σ_N_X__* = *s_N_X__* and *σ_N_Y__* = *s_N_Y__*. The correlation between the new signal, *N_c_*, and *S_X_* and *S_Y_* determines the specific combination of change in variance and correlation produced:

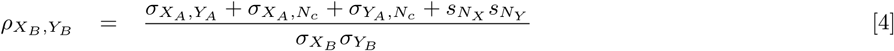

The range of correlations potentially explained by additions of a common signal will be smaller than the range that could be explained by the general ASC class. Typically, additions of common signal will produce increases in correlation, while additions of uncorrelated signal reduce correlation.

**Fig. 2.**
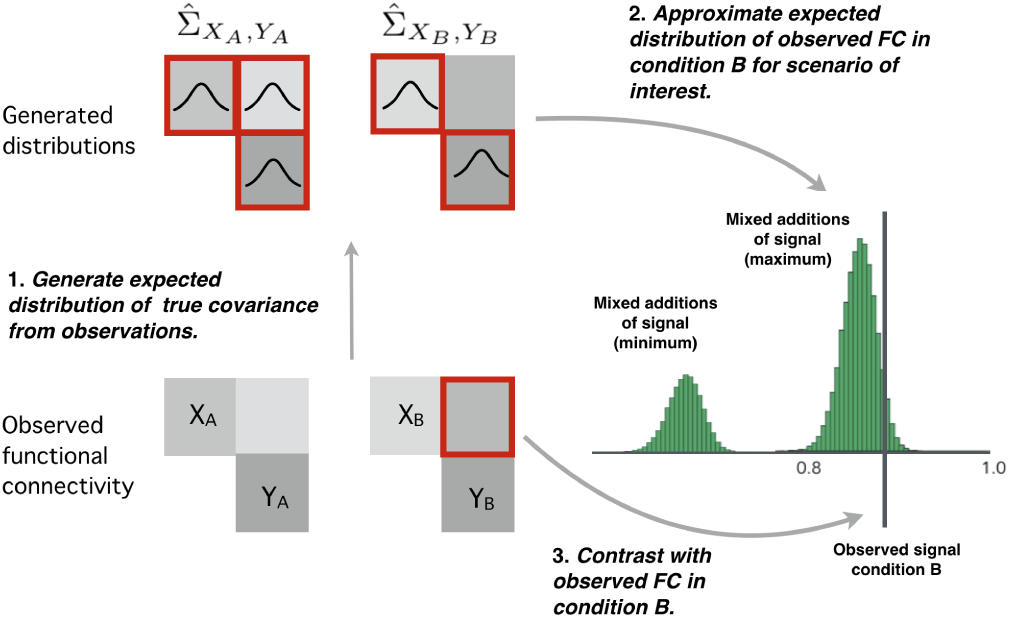
The Monte Carlo procedure used for inference on changes in functional connectivity. 1. From the observed covariances a distribution of potential underlying true covariances is generated, using a Wishart distribution and rejection sampling. 2. From these samples, the expected distribution of observed correlation for different additive signal change scenarios can be calculated (green histogram). These distributions can be compared to the observed correlation in condition *B*, which can be used to test the hypothesis that the observed data is explained by a given scenario. Note that addition of common and mixed signals can produce a range of changes in correlation, so distributions for minimum and maximum correlations are calculated.

### Statistical Inference

FC analyses typically make inferences regarding whether observed correlations indicate significant differences across conditions. The ASC model permits additional inferences regarding the possible nature of the signal changes underlying these changes. We take a Monte Carlo (MC)-based null-hypothesis approach, assessing the likelihood that additive signal changes, and the subclasses defined above, would produce the observed changes in correlation and variance. This approach takes into account the imprecision of measured covariances by sampling from a distribution of underlying putative true covariances (Fig.2). We first sample from the distribution of possible underlying covariances of the observed signals (Σ_*X_A_*,*Y_A_*_, Σ_*X_B_*,*Y_B_*_) by rejection sampling using inverse Wishart distributions (see Materials and Methods, *Inference Procedures*). This assumes a Gaussian generative signal and a flat prior. Appropriate degrees of freedom (dof) for the autocorrelated time fMRI series is estimated by fitting AR signal models to the signal (Fig.2).

To test for whether observed changes in covariance, *Q_A_*, *Q_B_*, could be produced by additions of uncorrelated signal (ASC subclass 1), we first sample from the distribution of underlying true covariances of condition *A*, the distribution of true variances in both conditions, given the observed signal. We then use Eqn. 4 and a Wishart distribution to generate samples of expected observed correlations in the second condition. This distribution is then compared to the observed correlation in the second condition, to determine whether uncorrelated signal changes would be likely to have produced this observation.

A similar approach can be used to determine whether an changes in covariance can be explained by an addition of a common signal component (ASC subclass 2), or by other additive signal changes. For these classes, specific variance changes can produce a variety of changes in correlation, depending on the nature of the introduced signals, with the change in correlation determined by the correlation of the additional signals with the existing signal. In these cases, we perform a search over the correlation between the putative additional signals and existing signals to determine the range of potential changes in correlation that could be produced. To perform inference, those signals identified as producing the maximum and minimum possible changes in correlation are used in the sampling approach described above, to produce separate distributions for the expected maximal and minimal value of correlations that can be explained in these scenarios (Fig.2).

In our demonstrative application of the above procedures, we assessed multiple connections within a network of brain regions. To present the results, we first identified connections showing a significant change in correlation (corrected the using False Discovery Rate, *α* = 0.05). These are presented in four connectivity charts, showing changes in correlations that: 1. can be explained by uncorrelated signal, 2. can be explained by common signal, 3. can be explained by other additive signal changes, or 4. cannot be explained solely by additive changes of signal. Membership of these classes was determined using the MC approach above with 2000 iterations.

### Model validation and simulations

*Basic simulations.* We first demonstrate the types of changes in signal that additive signal can produce using simulations. Fig.3 shows the effects of additive signal changes on the observed correlation of two nodes with initial correlation of 0.58. The white histogram shows the distribution of the observed correlation in this initial condition. We simulated the effects of additions of signal producing 20% increases in the standard deviation of both nodes. The red histogram shows the distribution of observed correlation of the second condition the additive signal was uncorrelated across nodes. On average, an addition of uncorrelated signal produced a drop in observed correlation to 0.40, with a 95% range of (0.35, 0.46). The blue histograms show the increases in observed correlation produced when a common signal component was added to both regions. Here, the extent of increase will depend on the correlation of the new signal, **N_*c*_** to the initial signals. The two histograms correspond to the effects of those common signals that produce the smallest and largest changes in correlation. Common signals producing a smallest change in correlation on average produced an observed correlation of 0.64, with a 95% minima of 0.61. Those common signals producing the largest increase in signal produce an average observed correlation of 0.71, with a 95% maxima of 0.73. Changes produced by the broader class of additive signals are represented by the pair of green histograms. This class can account for a wider range of changes in correlation associated with the 20% increase in standard deviation. Here, the minimum correlation is distributed around 0.02, while the mean maximum correlation is 0.86, with an overall 95% interval of (-0.09, 0.89). Mixtures of additive signals may produce correlations lower than that produced by increases in uncorrelated signal as the signals added to the two nodes can be negatively correlated with each other. Supporting results shows how initial correlation alters the effects of different additive signals (Figs. 2).

**Fig. 3.**
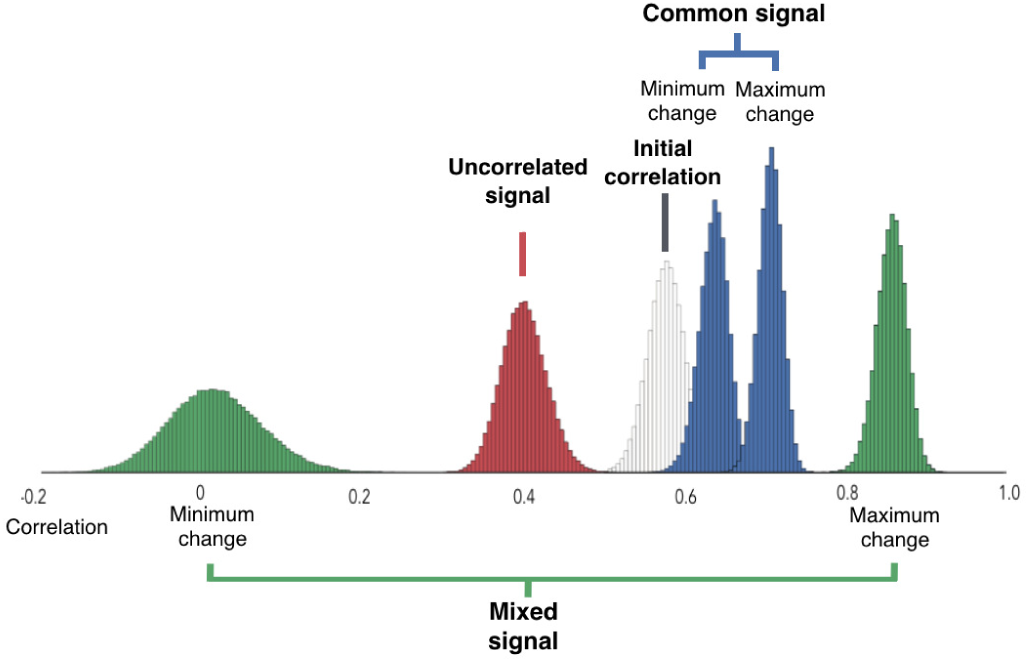
The effects on correlation of additive signals producing a 20% variance change. The plot reflects a scenario where both regions increase in variance by 20%, with an initial correlation of 0.58. The white histogram reflects the distribution of observed correlations in condition *A*. The red histogram represents the expected distribution of correlation in condition *B*, if the observed 20% change in variance was associated with uncorrelated signals. The blue histograms reflect the distributions of maximum and minimum changes in correlation if variance changes were due to a common additive signal component. Finally, the green histograms reflect the distributions of maximum and minimum changes in correlation when variance changes are due to any additive additions of signal.

*Inference on simulated network changes.*
Fig.4 shows results from an analysis of a simulated scenario of the addition of a common signal into certain nodes of a 10 node network, using the Monte Carlo inference procedure. Here, additions of a common signal into three nodes produced 20% changes in variance. The additive analysis procedure correctly identified nodes sharing a common increase in signal. Additionally, it correctly identified decorrelations of these nodes with other nodes not receiving the signal as changes consistent with increases in uncorrelated signal. This occurred only for connections with some initial correlation. The lower plots indicate the specific changes in correlation for those connections showing a change in correlation, and the range of changes that could be explained by ASC. Note that for these relatively large variance changes, a broad range of changes in correlation could be explained by additive changes in signal.

**Fig. 4.**
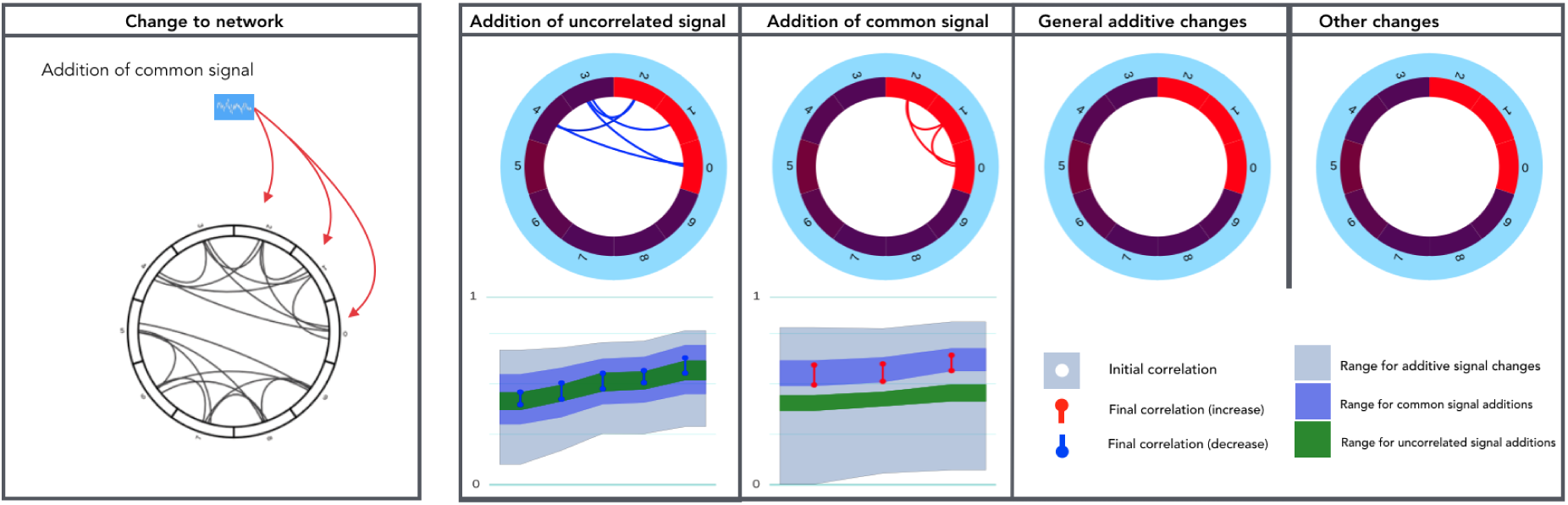
Example analysis of simulated changes in brain networks. Grey connections in the first column indicate nodes with a positive correlation (between 0.3 and 0.7) in the initial condition, and red arrows point into nodes to indicate that additional signal was injected in the second condition, producing an increase in variance of 20%. Here, a common stochastic process was added to nodes 1-3, uncorrelated with existing signal. The remaining columns identify connections that were detected as showing significant changes in correlation across conditions, FDR corrected (*α* = 0.05). Red connections indicate increases in correlation, blue decreases. Node colors represent change in variance. Each circle plot shows connections that can be explain by a particular class of additive change, with the final column showing remaining connections with significant change in correlation. The underlying plots show the distribution of correlation changes for the shown connections. White dot - correlation in initial condition. Blue/red dot - correlation in second condition. Colour fill - range of correlation in second condition that could be explained by additive signal change. The increased correlations between the three nodes receiving the introduced signal are identified as indicative of increases in common signal. Decreases in correlation between these nodes and other correlated nodes are identified as indicative of increases in uncorrelated signal.

### Analysing functional connectivity changes in experimental FMRI dataset

We assessed the extent to which the analysis approach informs functional connectivity analyses by applying it to an FMRI study of functional connectivity changes across resting and steady-state motor and visual task conditions (see **Materials and Methods**, Supp Fig. 1). Fig.5 presents the circle-plot results from the additive signal analysis of contrasts of a standard resting state and states in which continuous, visual stimulation (A) and finger tapping (B) was occurring. For clarity, the ROI nodes are restricted to consist of regions relevant to the visual and motor tasks, identified in a prior localisation procedure. Further contrasts involving combined visual and motor conditions are shown in Fig. 6. Seed-based mapping showed that the continuous task conditions produced substantial changes in both correlation and variance (Supp Fig. 3).

**Fig. 5.**
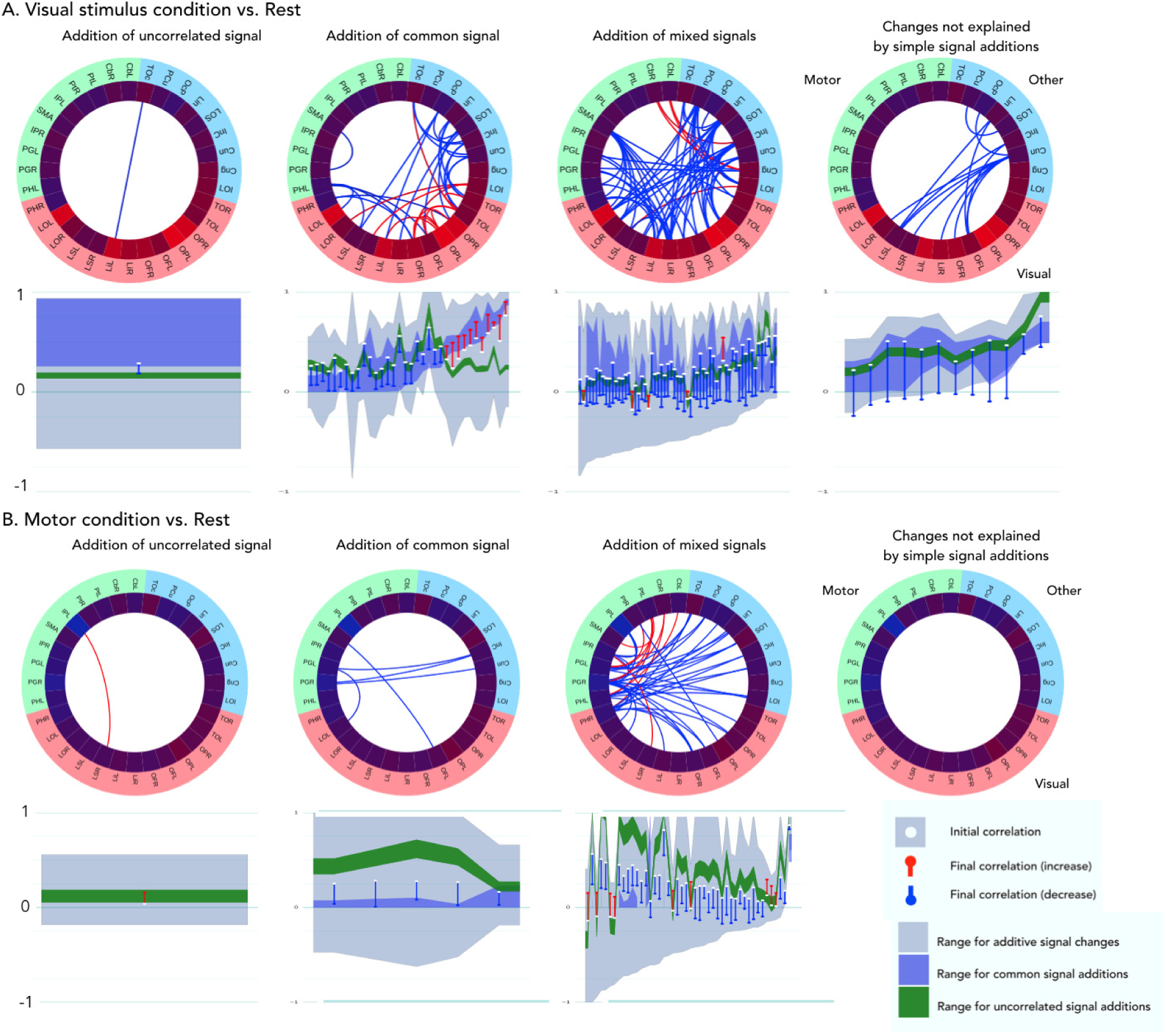
ASC analysis of FC changes between rest and visual stimulation (A), and rest and a motor condition (finger tapping) (B). Plot organisation is described in Fig. 4, region labels in Table 1 (Supporting Information). The majority of significant changes in correlation can be explained by additive changes in signal. The visual condition produced increases in variance in visual regions relative to rest, and was associated with increases in correlation between visual nodes that could be explained by increases in common signal or other additive signal changes. Decorrelation with other regions could also be explained by additive changes. The motor condition produced modest reductions in variance in motor regions, which were nevertheless enough for additive changes to explain the majority of changes in correlation, including decreases between cortical motor regions, and increases in correlation of cortical motor regions with subcortical motor regions.

**Table 1.** Region label key

|  |  |
| --- | --- |
| Cng | Cingulate Cortex |
| Cun | Cuneus Cortex |
| InC | IntraCalcarine Sulcus |
| SMA | SMA |
| LOI | Lateral Occipital Cortex (Inferior) |
| LOS | Lateral Occipital Cortex (Superior) |
| Lin | Lingual Gyrus |
| OcP | Occipital Pole |
| PCu | Precuneus Cortex |
| TOc | TempOcc |
| CbL | CbL Cerebellum (Left) |
| CbR | CbR Cerebellum (Right) |
| IPL | Inferior Precentral Gyrus (Left) |
| IPR | Inferior Precentral Gyrus (Right) |
| PtL | PutL Putamen (Left) |
| PtR | PutR Putamen (Right) |
| SMA | SMA |
| PGL | PostGL Postcentral Gyris (Left) |
| PGR | PostGR Postcentral Gyris (Right) |
| PHL | Premotor Cortex Hand area (Left) |
| PHR | Premotor Cortex Hand area (Right) |
| LOL | LatOccSL Lateral Occipital Cortex - inferior div. (Left) |
| LOR | LatOccSR Lateral Occipital Cortex - inferior div. (Right) |
| LSL | LatOccSL Lateral Occipital Cortex - superior div. (Left) |
| LSR | LatOccSR Lateral Occipital Cortex - superior div. (Right) |
| LiL | Lingual Gyrus (Left) |
| LiR | Lingual Gyrus (Right) |
| OFR | Occipital: Fusiform Gyrus (Right) |
| OFL | Occipital: Fusiform Gyrus (Left) |
| OPL | Occipital Pole (Left) |
| OPR | Occipital Pole (Right) |
| TOL | Temporal Occipital (Left) |
| TOR | Temporal Occipital (Right) |

**Fig. 6.**
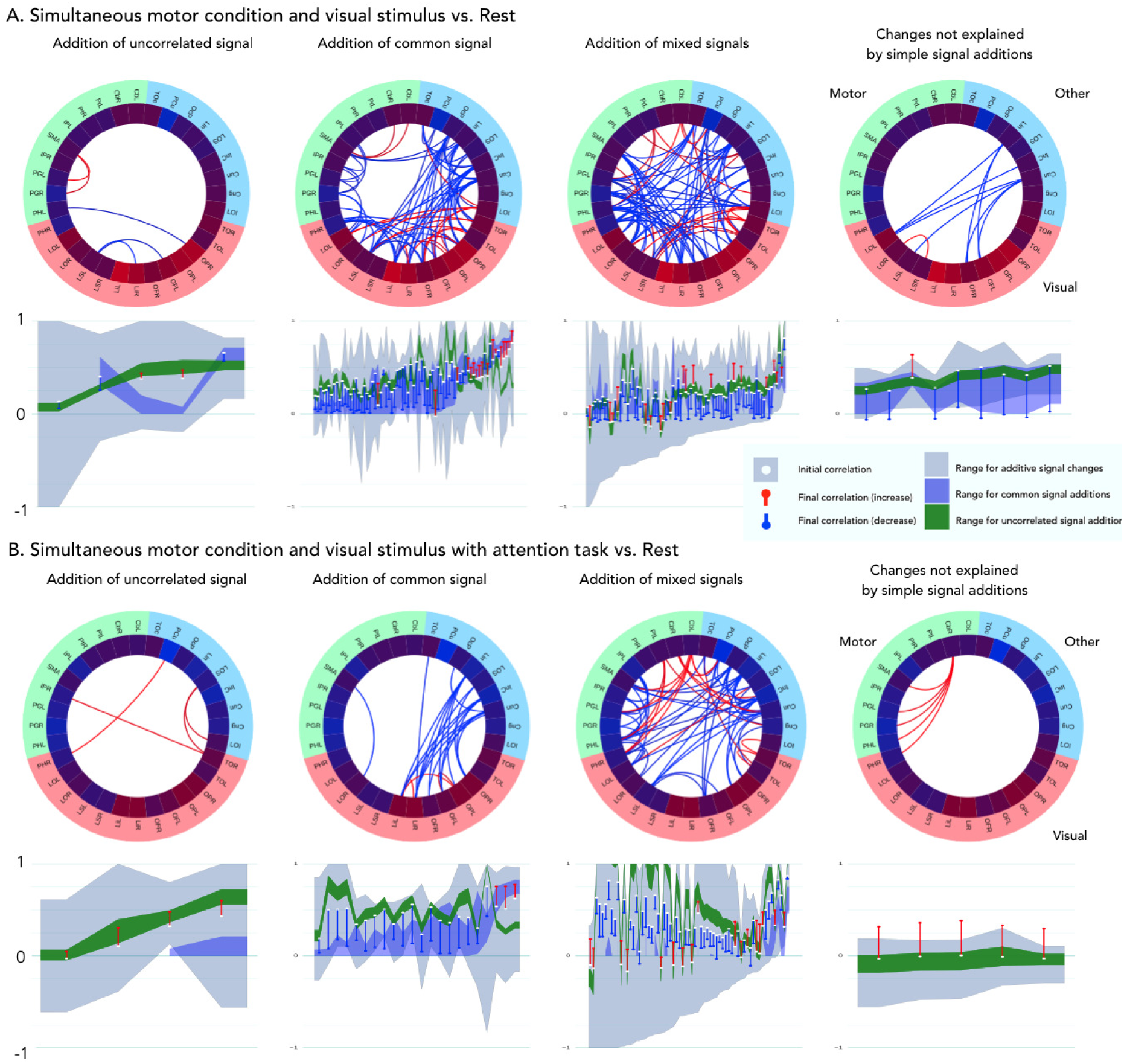
Connectivity changes between combined finger tapping and visual stimulus condition and rest, and a more attention demanding combined task and rest. Plot organisation is described in Fig. 4

Overall, many node-pairs showed differences in correlations across the rest and the steady-state conditions, many of which were lower during the active conditions compared to rest. Many of these changes could be explained by additions of a single common shared signal (column 2), or other additive signal changes (column 3). The pattern of these changes showed consistencies for conditions involving the same task types, with the ASC providing considerable insight into the possible nature of the signal changes under lying the different task states.

Visual stimulation (Fig. 5A) produced strong increases in variance within visual regions. This was associated with increases in correlation between certain visual brain regions that could be explained by increases in a common shared signal component. A larger number of reductions in correlation could also be explained by the increases in signal in visual regions. Here, this change reduced correlations indicating that the new signal in the visual regions was weakly correlated with activity in the non-visual regions. These changes were sometimes classified as being explained by changes in common signal, which can occur if the change in variance in one node is small or negative. A number of reductions in correlation between activated visual regions and regions not activated could not be fully explained by additive signal changes, suggesting, perhaps, a combination of increases and decreases of signals in these nodes.

The finger tapping task (Fig. 5B) produced modest reductions in variance in motor-specialised regions, along with a reduction in correlations between these regions and other regions. These changes could be explained by either reductions in common signal, or general additive changes in signal (i.e. additional common or partly shared signal in the rest condition compared to finger tapping). In many cases, correlation reduced from around 0.2 to close to 0. The motor cortical regions also showed some increases in correlation with specific brain regions associated with motor control, namely putamen and cerebellum. These changes could be explained by additive changes, corresponding here to the motor cortical nodes having additional uncorrelated signal at rest that reduced in the motor condition.

The visual-motor condition (Fig. 6A) produced a pattern of extensive effects with correspondences to the two single-task conditions, with visual and motor cortical regions showing similar increases and decreases in variance. The final condition, which required subjects to attend to specific changes in the movie, demonstrated similar changes to the similar combined task ( Fig. 6B), produced more extensive reductions in signal variance, particularly in default mode regions. Increases in variance visual regions diminished, possibly due to the broad reductions in variance counteracting the local visual-stimulus related increases in variance. Increases in correlations between motor cortical regions and cerebellum, which were near zero at rest, could not be explained by additive changes in signal.

## Discussion

Considered on its own, correlation provides an ambiguous characterisation of how the relationships between signals change. We have presented an approach identifying additive signal change (ASC), which enhances correlation-based analyses, identifying changes that can be explained by simple additions of signal, or changes in amplitude of signal components. It provides a characterisation that can be used almost anywhere correlation is used, requiring few assumptions and limited computation. The classes of the ASC analysis correspond to natural and distinct scenarios that can be expected to appear in many scenarios. Our results on FMRI data confirm this, suggesting that many changes in FMRI correlation can be explained by additive changes in signal. This has important implications for the interpretation of FC analyses, where changes in correlation are often broadly interpreted as indicative of changes in neural coupling of brain regions.

### Interpretation of functional connectivity with ASC

The ASC analysis approach clarified how functional activity and the relationships between nodes varied across the five conditions of the FMRI dataset. A majority of the observed changes in correlation were associated with changes in variance in one or both nodes that were substantial enough to explain the correlation changes in terms of additive signal changes. For example, the increases in correlation between visual regions during visual stimulation could be explained by an addition of common signal corresponding to the observed increases in variance. These putative additions of signal could also explain the decorrelations between visual regions and regions that did not show a similar increase in variance. The ASC analysis provided insight into the striking differences in effects of motor and visual tasks on FC, where these tasks had opposite effects on the variance of activity in activated brain networks. Changes in measured FC will be sensitive to differences in the stability and amplitude of activity across states, which may vary for a variety of reasons. The ASC analysis identified some cases where changes in correlation could not be explained by additions of signal. For example, the motor task produced increases in correlation between cerebellum and SMA, from around zero, that could not be explained entirely by additive signal changes. Reviewing ASC results across a variety of connections and conditions provides further insight into the nature of FC changes. For example, the visuo-motor attention task appeared to broadly dampen fluctuations, including in regions where visual stimulation had otherwise increased the amplitude of fluctuations. Visual regions showed correspondingly smaller increases in correlation. Changes induced by head-motion will typically be additive, and show particular spatial patterns (17). These results make it clear that correlation changes on their own may provide poor insight into the complex array of changes in underlying changes in FC. Evidence of changes in intrinsic coupling and decoupling between nodes can be detected, but must be carefully distinguished from changes that might be explained by additive changes in the amplitude of signal components.

### Relevance to other methods

Similar challenges for interpretation apply when covariance or regression co-efficients are studied in isolation. In FMRI, functional connectivity analysis approaches such as psycho-physiological interactions (PPI) (18, 19) and dual regression (20) estimate group level statistics of regression coefficients of signals derived from preselected nodes or networks applied to individual voxels. The sources for observed changes in these analyses remain ambiguous unless variance is explicitly characterised. ASC analysis could be used in a seed-based mapping analysis, identifying all voxels showing a particular class of correlation change with a given seed. The approach described here has analogues in the frequency domain, where additive changes signal components predictably alter coherence (3).

### Applicability to resting state studies

While the results presented here reflect modulations due to continuous induced task-driven activity, the methods and conclusions are relevant to studies of pure rest conditions, such as differences across patient groups. Along with the many studies reporting differences in resting state FC across different subject groups, numerous studies have reported differences in the amplitude of fluctuations (15, 16). However, variance changes may be more challenging to track in multi-session and parallel group studies, as there will be additional session-to-session and subject-to-subject sources of uncontrolled variability. Careful denoising and normalisation of signal amplitudes will increase sensitivity.

### Relation to causal modeling

The ASC analysis will provide a useful guide for designing and interpreting effective connectivity modelling strategies such as DCM (10) where model estimation will be strongly influenced by the covariance structure. In neuroscience, causal modelling approaches are often focused on characterising the structure of directed connections within smaller sized networks. In this context, additive signals could be produced by increases in incoming connections to a node, or indirect effects associated with changes in connectivities of nodes with indirect connectivity to the node of interest. A prior ASC analysis could be used to identify patterns of functional connectivity likely to influence causal modelling, and guide ROI selection and other model choices. When interpreting causal model fits it will be useful to cross-check identified changes in coupling with their implied additions of signal to specific nodes. ASC mapping analysis could also be used to map regions not included in the causal model that show similar overall patterns of covariance change, which will indicate the spatial distribution of brain activity that may be associated with a particular causal role.

### Further work

A number of improvements and extensions could be made to the ASC protocol. The present analyses is either performed on individual data samples or concatenates across samples (e.g. subjects). It may be possible to devise a random effects or other hierarchical modelling approach that still provides useful categorisation of changes (21, 22). ASC is straightforward to apply to partial correlation matrices. Here, the putative shared variance components will reflect variance unique to the pair of regions under investigation. Related extensions would be to extend the shared component model to characterise three or more regions simultaneously, identifying signals that are shared across subsets of regions. Another strategy to provide finer dissection of variance components could be to simultaneously model multiple conditions simultaneously.

We have presented an additive signal change model linking correlation and variance that can substantially enhance the description of functional connectivity. The additional information provided by classifying correlation changes into those that can be explained by specific simple changes in signal, and those that are suggestive of coupling changes, provides a less ambiguous characterisation of functional connectivity that may make it easier to share and assess studies of functional and effective connectivity.

## Materials and Methods

### Experimental protocol and acquisition

Sixteen healthy volunteers were scanned under five separate five-minute steady-state conditions, with no baseline epochs: rest (eyes open), visual only, motor only, simultaneous (but independent) visual and motor tasks, and a combined condition involving a visually-cued motor task (Supp Fig. 1). The visual conditions consisted of videos of colourful abstract shapes in motion. The motor conditions involved continuous and monotonic sequential finger tapping against the thumb, using the right hand. The combined motor conditions combined the visual stimulus with finger tapping. In the visually-cued motor task subjects where instructed to change tapping direction when they saw an irregularly appearing cue, which were present in all visual conditions. These conditions were designed to induce robust changes in functional connectivity within well defined networks. An additional block-design task-activation localiser FMRI scan was performed under the same conditions to enable the identification of brain regions changing in average activation levels during these conditions. This scan used pseudo-randomised 30-second block intervals separated by 30-second baseline periods. Data was acquired in a Siemens 3T scanner, using a 32-channel coil and a high-resolution (2*mm*^3^) fast (TR = 1.3s) multiband (factor 6) whole-brain acquisition (23, 24). Scans were five minutes (230 time points).

### Correlation change produced by additive changes in signals

Here the aim is to find the limits of:

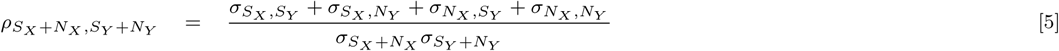

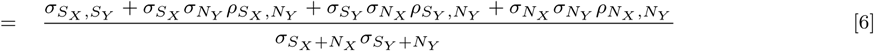

with the constraints:

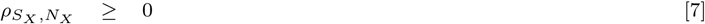

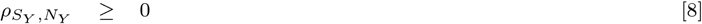

Expressions for *σ_N_X__* and *σ_N_Y__* can be obtained:

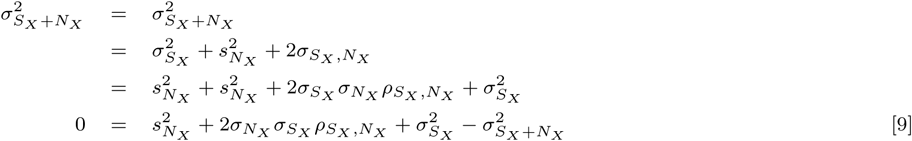

Solving the quadratic gives values for *σ_N_X__*:

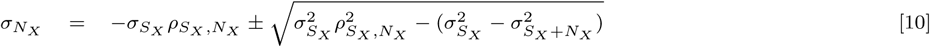

*σ_N_Y__* can be obtained similarly.

We optimise Eqn. 6 with a search over values of *ρ_S_X__*,*_N_x__*, *ρ_S_Y__*,*_N_Y__*. From these values and the measured *ρ_S_X__*,*_S_Y__* we can identify maximum and minimum values for *ρ_S_X__*,*_N_Y__*, *ρ_S_X__*,*_N_X__* and *ρ_N_X__*,*_N_Y__* using the Cauchy-Schwarz inequality applied to correlations:

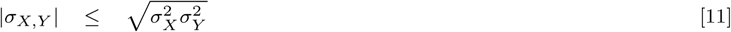

Obtaining these results involves the decomposition of processes into orthogonal components, where, 
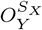
 defines the component of *X* that is orthogonal to *Y*. We have:

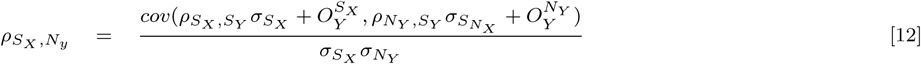

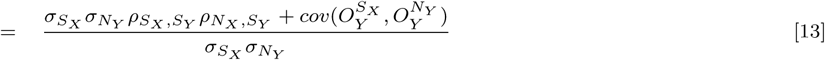

With 
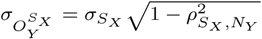
 we apply applying the Cauchy-Schwarz inequality to the final term:

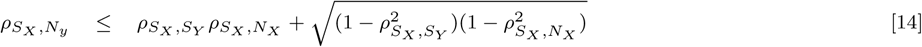

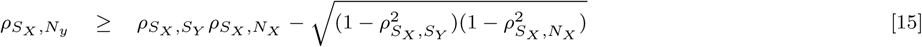

### Correlation change with the addition of a common signal

Limits for the addition of a common signal can be calculated similarly, where we are now optimising:

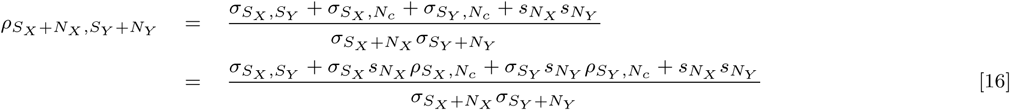

### Scenarios where increases in variance occur in different conditions

The formulations described can be transfer to scenarios where additions of signal occur in different conditions in the two nodes. Here, we the flip constraint on correlation for one node of Equations 7 in formulation above, to obtain *ρ_S_X__*_+_*_N_X__*,_−_*_N_X__* > 0

### Monte-Carlo Inference

To sample the possible distribution of covariance matrices underlying the observations, we assume that their distribution is equal to that of covariance matrices of normally distributed correlated variables sharing the same degrees of freedom (*n*) (accounting for the autocorrelation in the FMRI data). A *p* × *p* covariance matrix Σ, of Gaussian variables with *n* degrees of freedom, will have a Wishart distribution, with a pdf:

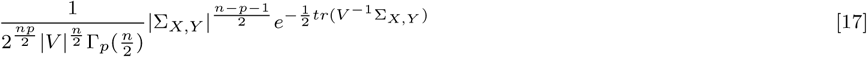

Where Γ_*p*_() is the multivariate gamma function:

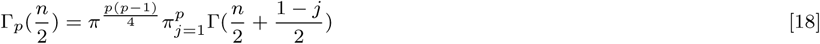

Using a flat empirical prior, we used rejection sampling to to obtain samples from the distribution of underlying covariance matrices A and B potentially producing the observed covariances, **Q_A_**, **Q_B_**. For each sample, limits on the range of correlations produced by additive, uncorrelated and common signals can calculated. Additional variability will occur in scenarios with a low initial correlation (or low degrees of freedom), where different samples will have the underlying initial correlation between signals vary in sign.

### FMRI Preprocessing

Data were analysed using FSL’s FEAT (25), the FSLNets network toolbox (4), and with Python, using SciKits-Learn (26) and the MNE package (27). Standard FEAT preprocessing was applied, including brain extraction (28), motion correction (29), and 2mm spatial smoothing. To reduce noise, we used our automated denoising tool, FIX (FMRIB’s ICA-based Xnoiseifier) (30), which removes artefactual signal components associated with head motion, physics artefacts, and non-neural physiological signals. FIX also integrations a regression of measured head motion parameters. Checks of image quality, head motion, and registration were performed for every scan. Data was high-pass filtered (0.005 Hz) to remove low-frequency drifts and other artefacts that would affect variance estimates. Variance maps and variance change maps were generated using the FSL tools FSLMaths and Randomise. Correlation maps were generated using FEAT and specialised code written in Python.

### Modelling of localiser scans

A block-design task activation scan was acquired to identify those regions showing task-related activation to the steady-state tasks investigated here. This involved a nine-minute scanning period that involved randomised 30-second blocks of the visual, motor and motor-visual tasks (total of 12 blocks). The motor-visual attention task was not acquired in this scan. Due to scanner limitations, this scan used a non-accelerated sequence, with a repetition time of 3.0s and 3mm cubic pixel resolution.

A multi-level GLM analyses of the localiser scan was used to identify regions that activated with one or more of the conditions, relative to rest (25). The block design time series of each of the three conditions was convolved with a gamma function, and included within a model also included the temporal derivatives of these time series, and motion parameters derived from motion correction. These analyses used temporal autocorrelation correction and outlier detection to ensure model validity (32, 33). Subject-level contrasts were defined for each of the three task conditions.

The resulting parameter maps were registered to standard anatomical space, via their high-resolution structural images using the FSL linear and non-linear registration tools FLIRT and FNIRT. The parameter maps were then combined at the group level in an F-Test to identify brain regions showing positive or negative responses to one or more of the conditions. The maps were transformed into Z-statistic images and thresholded (correcting for multiple comparisons) using FSL FEAT with a cluster-based thresholding approach with cluster-level significance level of p<0.05.

### Regions of interest

33 task-specific regions of interest were defined by identifying clusters of voxels showing significant responses to the combined task conditions. Their delineation was achieved by intersecting active clusters with regions of the Harvard-Cortical Atlas (https://www.fmrib.ox.ac.uk/fsl). The analyses comprised both whole-brain mapping analyses and ROI-based characterisation of connectivity maps. To increase specificity, we targeted regions activated or deactivated in the task localiser scans. To identify task-related regions, we delineated regions identified as significantly activated in the localiser task-activation analyses (using an F-test across conditions). To separate nodes, these regions were intersected with the Harvard-Oxford parcellation of the cortex (https://www.fmrib.ox.ac.uk/fsl).

To generate parcellations, group-level MELODIC output was matched to 10 canonical RSNs as defined by (Smith et al., 2009) via spatial cross-correlation methods as described above. Thresholds for the ICA maps were set to Z>3. Then the spatial overlap for each FCN was compared to probability structures in the Harvard-Oxford Cortical (probability threshold was set to p>.5) and Subcortical Atlases, as well as with structures in the Cerebellar Atlas. No nodes within networks overlapped. This procedure resulted in each network being parcelled into atlas structures. Additionally, activation maps were thresholded at Z>2 and parcelled using the same algorithm. Average time series for each parcel and for whole ICNs were extracted and correlated, and covaried with one another to generate two sets of correlation and covariance matrices, one with respect to entire RSNs and one with respect to parcelled RSNs, for each of the four task conditions.

In order to explore the interaction between local activation, ICNs, and FC, activation maps were parcellated into ROIs using an automated technique reliant on the Juelich and Harvard-Oxford probability atlases as described in Section 3.2.5 (Parcellation (2)). The purpose of this analysis was to better understand how activated regions change in connectivity in relation to deactivated regions. The other goal of the analysis was to examine the difference between activation which spatially coincides with ICNs and activation which does not.

Source code for performing these analyses will be made available for download: git.fmrib.ox.ac.uk/eduff/ampconn Source data will be submitted to Neurovault: https://www.neurovault.org.

## ACKNOWLEDGMENTS

We are very grateful for the multiband pulse sequence and reconstruction algorithms from the Center for Magnetic Resonance Research, University of Minnesota. We also thank Sasidhar Madugula, Roser Sala, Hayriye Cagnan, Thomas Nichols, Saad Jbabdi, Gerard Ridgeway and Úrsula Perez Ramirez for their contributions to processing the data, developing the codebase and reviewing the manuscript.

---

**Supplementary Figure 1.**
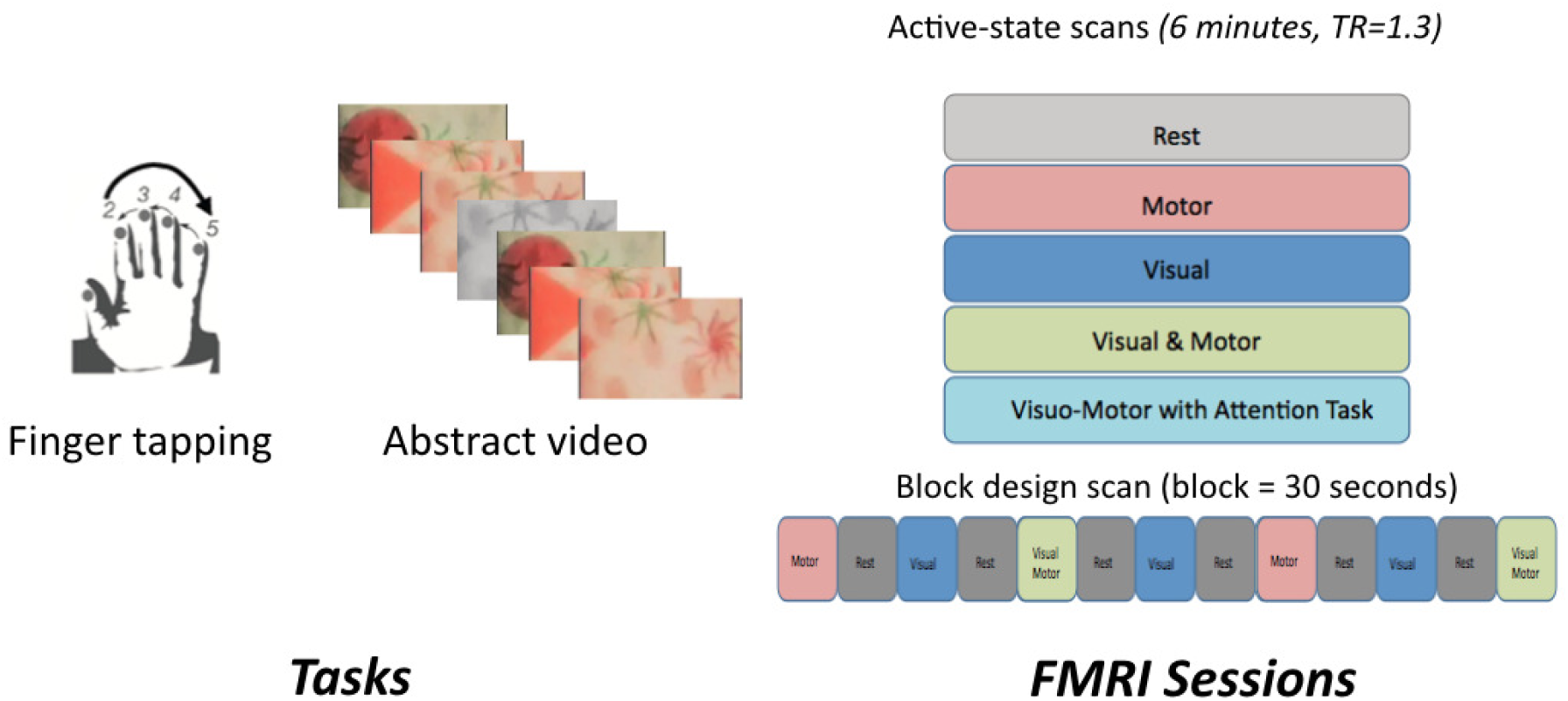
Validation experiment. A. Steady state tasks involved one or both of continuous fingertapping and viewing of a rapidly changing random images. The fingertapping had a consistent order, which was periodically reversed. B. Five 6-minute steady state conditions were used. In Visual & Motor condition subjects simply performed the motor task while viewing the video. In the attention task condition subjects were cued to change direction when specific visual cues were observed in the video.

**Supplementary Figure 2.**
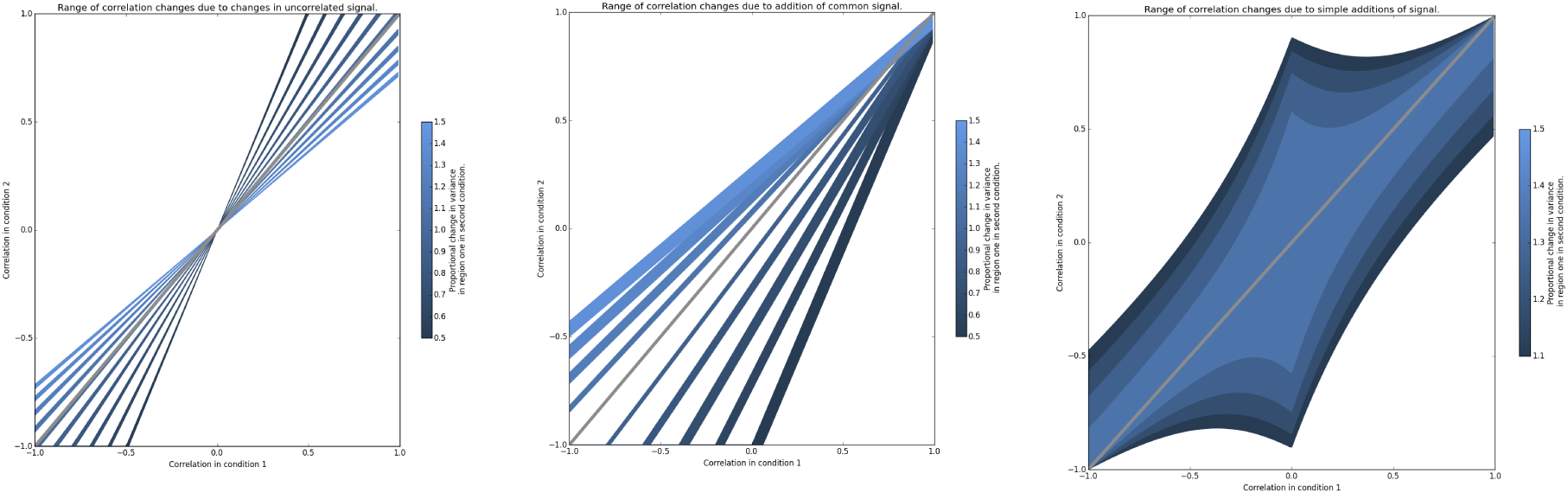
The effects on correlation of changes in signal component magnitudes producing certain variance changes, without observational uncertainty. The plots reflect scenarios where both regions change in variance (measured as standard deviation). Colour fills indicate the range of levels of correlation in the second condition that could be explaine.d by a specific change in standard deviation, if it reflects some simple combination or shared and unshared components. Column **a.** Potential changes associated with changes in uncorrelated signal. Column **b.** Potential changes associated with changes in a single signal component. Column **c.** Changes associated with a mixture of signal components changing in amplitude (only displayed for increases in variance).

**Supplementary Figure 3.**
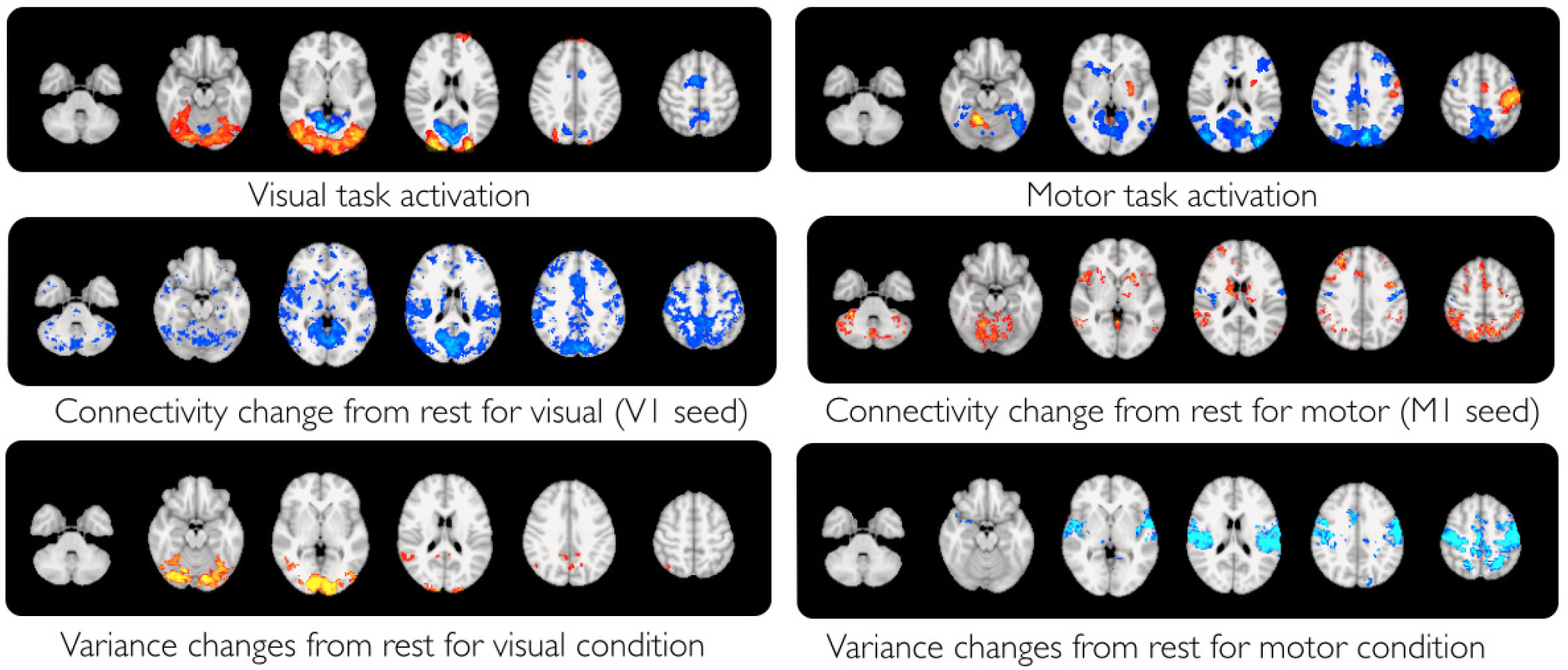
Changes in activation levels and variance across steady-state conditions. Maps show regions with significant increases (red) and decreases (blue) in standard deviation for stimulus conditions compared to rest. Histograms show the distribution of proportional changes in visual and motor networks.

